# Liver cells and PBMC secrete Tumor-suppressive Plasma Extracellular Vesicles in Melanoma patients

**DOI:** 10.1101/809301

**Authors:** Jung-Hyun Lee, Martin Eberhardt, Katja Blume, Julio Vera, Andreas S. Baur

## Abstract

Before and after surgery melanoma patients harbor elevated levels of extracellular vesicles in plasma (pEV), but their cellular origin is obscure. Here we suggest that these pEV are secreted in part by tumor cells, but particularly by liver and peripheral blood mononuclear cells (PBMC), which strongly suppressed tumor cell activity. As the cellular origin of pEV is difficult to determine, we mimicked the interaction of tumor cells with liver cells and PBMC in vitro, and compared newly secreted EV-associated miRNAs and protein factors with those detected in melanoma patient’s pEV. The results identified factors that could be associated either with tumor cell activity or the counteracting immune system and liver cells. Notably, the presence/absence of these factors correlated with the clinical stage and tumor relapse. Our study provides new insights into the innate immune defense against tumor cells and implies that residual tumor cells may be more active than previously thought.

**Summary blurb:** Plasma extracellular vesicles (pEV) in melanoma patients are a mix of cancer cell-suppressive vesicles from liver cells and PBMC, but derive also from residual cancer cells.

## Introduction

Human plasma is considered to harbor a high concentration and mixture of plasma EV (pEV) of different cellular origin. However, confirmed or accepted numbers for pEV, or specific subpopulations thereof, are lacking. This is due to technical difficulties in identifying and discriminating vesicles. In addition, the great variability of pEV with respect to size, surface marker composition, cellular and subcellular origin further complicates this task. Nevertheless it was suggested that a significant portion of pEV are secreted by platelets, namely up to 10^7^/ml plasma, as determined by flow cytometry (1). Notably, these pEV were found to have multiple effects on target cells in homeostasis and disease (2).

There is accumulating evidence suggesting that cancer cell-derived pEV harbor specific biomarkers (3, 4). However, there is insufficient knowledge on their relative blood concentration in different stages of the disease, particularly after complete (R0) tumor surgery. Complicating the situation, cancer cells may stimulate host cells to secrete additional pEV populations. Tumor cells and their secreted EV may stimulate immune cells and/or the tumor microenvironment, for tumor-suppressive or tumor promoting effects (5, 6). For example, senescent tumor cells secrete the senescence-associated phenotype (SAPS), consisting of a whole array of pro-inflammatory soluble and insoluble factors, able to stimulate fibroblasts of the tumor microenvironment (7). Notably, the SASP may also be secreted through EV (8).

After purification by differential centrifugation and sucrose gradient, we determined that healthy individuals harbor around 10^9^ EV per milliliter plasma. This vesicle concentration increased between 4-20fold in disease conditions, like malignant melanoma and HIV infection, and remained at higher levels after primary tumor resection or anti-retroviral therapy (9, 10). The relative pEV increase was corroborated by measuring vesicle-associated micro-RNA levels and protein concentrations in EV sucrose gradient fractions. However, it remained unclear from which cell population(s) these increased pEV levels was/were secreted.

Since we found that elevated pEV after R0 surgery were tumor-suppressive (9), and since pEV have a rather short life span of about 10-30 minutes (11, 12), a sizable and active cell population or organ had to be assumed, serving as their cellular source. On the other hand, a tumor cell-derived secretion seemed possible, although the in general low concentration of circulating or disseminated tumor cells (CTC/DTC) (13), as well as their assumed tumor promoting function, suggested a minor contribution to the increased pEV levels after surgery.

In view of the difficulties to determine the cellular origin of pEV, we analyzed the EV content from co-cultures of PBMC and liver cells with tumor cells in vitro. We reasoned that this mimicked the interaction of these cells in vivo, producing unique factor patterns that would be detected in patient’s pEV, if our assumption was correct. Our results supported this hypothesis and pointed to a strong vesicular secretory activity of liver cells and PBMC. To our surprise, we also registered a prominent activity of residual tumor cells. Our study provides unexpected insights into the interaction of tumor cells with the host that could be exploited for screening and monitoring of cancer.

## Results

### Hepatocytes are a possible source of pEV

Since CTC/DTC are usually found at very low frequency (10^−5^-10^−6^) in bone marrow and lymph nodes (Klein, 2000), we assumed that a major fraction of pEV in post-surgery melanoma patients were of non-tumor origin, as for example from the innate immune system. To test this assumption, we compared miRNA profiles in EV secreted from primary immune cells with those of patient’s pEV and healthy controls. For this analysis the same patient data set was used as described in our recent study (9). In that study patients had been subdivided into those with a high risk (**HR**) and a low risk (**LR**) for tumor relapse, and those bearing a tumor (**T**).

We employed correspondence at the top (CAT) plots (14), comparing EV/pEV miRNA profiles that were ranked by relative miRNA concentrations, assuming that profiles from the same cell linage have a similar ranking. A pairwise comparison of the 150 highest-ranked miRNAs, revealed that EV-derived miRNA profiles from primary immune cells (macrophages, immature and mature DC) had only a low correspondence with patient’s pEV-derived profiles (30-40%), and no noticeable difference between patients and healthy controls was observed (**Fig 1A**).

**Figure 1:**
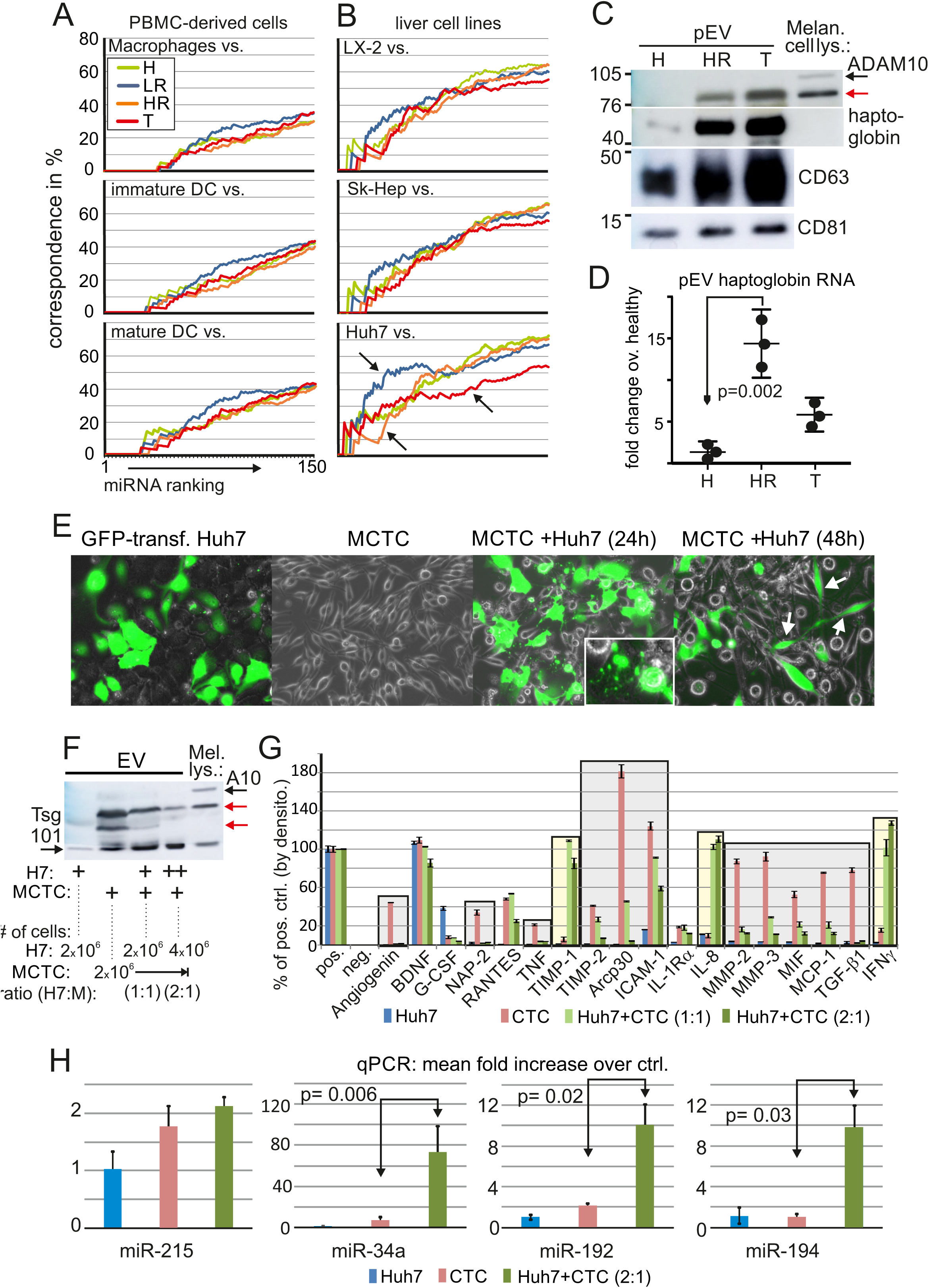
Hepatocytes are a possible source of melanoma-suppressive Pev. **A-B** Relative similarity of pEV miRNA profiles with cell-derived EV profiles using correspondence at the top (CAT) plots. **A** Pairwise similarity (correspondence in %) of 150 ranked miRNAs from EVs derived from myeloid cells (mature DC, immature DC, macrophages) and **B** liver cells (Huh7, SkHep-1, LX-2), which were compared with pEV-derived miRNAs from healthy individuals (H) and melanoma patients (LR-, HR- and T-patients; see text for explanation). Arrows indicate deviation of correspondence from control (Healthy). **C** Presence of haptoglobin in melanoma pEV. Plasma EV from one healthy individual and one HR and T-patient were gradient purified and blotted for the indicated pEV markers and haptoglobin as indicated. A melanoma cell lysate served as control for ADAM10. **D** Relative amount of haptoglobin mRNA in melanoma pEV compared to healthy donor pEV. Error bars represent SDM based on analyses of pEV from 3 different donors. **E** Images of GFP-transfected Huh7 and melanoma MCTC cells cultured separately or co-cultered over 48h at a ratio of 1:1. **F** Assessment of ADAM10 (A10) and Tsg101 (EV marker) by Western blot in EV lysates derived from cultures and co-cultures of Huh7 and CTC cells (90 ml supernatant), cultured in different ratios (1:1; 2:1). M= MCTC cells. H7= Huh7 cells. The overall cell number of MCTC cells remained the same (5×10^5^), the number of Huh7 cells increased to 10^6^. One lane represents the amount of EV purified from 15 ml supernatant. **G** Quantification of indicated factors assessed by protein array (raw data in **Fig S1B**) in EV secreted by cell cultures described in (F) (purified from equal culture volume). Raw data were quantified by densitometry and calculated in % from the positive controls (pos.) on each blot. Co-culture of Huh7 and MCTC cells were done with different cell ratios. Gray and yellow boxes are described in the text. Error bars represent SDM based on four values of two experiments (**Fig S1B**). **H** Quantitative PCR analysis on indicated miRNAs in EV obtained from cell culture supernatants described in (E). Bar diagrams depict the average fold-increase over an internal control miRNA of the commercial assay. Error bars represent the SDM of triplicate PCR runs, One representative experiment out of three is presented.

Recently we found evidence that pEV in HIV patients are secreted in part by the liver (15). We therefore included EV miRNA profiles derived from 3 liver cell lines into our analysis, representing parenchymal (Huh7), hepatic stellate (LX-2) and liver sinus endothelial cells (SK-Hep). Using CAT plot analysis, liver miRNA profiles revealed a higher overall correspondence with patient’s pEV and also healthy controls (60-70%), implying that at least a portion of pEV derived from liver cells. (**Fig 1B)**. Notably, when compared with hepatocyte (Huh7)-derived EV profiles, melanoma patient pEV miRNA profiles differed from healthy controls (**Fig 1B**, arrows). This hinted at hepatocytes as one possible source of cancer-induced pEV.

As these results were merely an indication, we analyzed patient pEV for liver-specific factors. We found that haptoglobin, a liver acute phase protein, was incorporated in significant amounts and we could also demonstrate an upregulation of its mRNA in pEV from patients bearing a tumor (T) and patients with a high risk (HR) for tumor relapse (**Fig 1C and D**). This supported our assumption that hepatocytes were at least one possible source for melanoma-induced pEV.

#### Upon tumor cell co-culture hepatocytes secrete tumor cell suppressive EV

For further clarification we decided to mimic an assumed interaction of hepatocytes with circulating tumor cells (CTC) in cell culture, asking whether this would induce hepatocyte EV secretion with tumor-suppressive properties, as implicated by the results of our previous study (9). Besides Huh7 hepatocytes we employed a melanoma CTC cell line (MCTC) established from peripheral blood of a patient in our department (see materials and methods for details).

Upon co-culture, both cells rapidly established contacts and after 24h GFP-transfected Huh7 cells had started a massive secretion activity, evidenced (1) by the appearance of green vesicular structures that were either secreted or transferred by protrusions (**Fig 1E**, panel 3, see insert) and (2) by an increase of particle/vesicle concentration in culture supernatants by more than 10fold over EV produced by the same cells without co-culture (**Fig S1A**). As a consequence, most CTC cells became GFP positive after 48h, likely due to vesicle uptake (**Fig 1E**, panel 4, see arrows).

To assess the consequences of this effect, EV were purified from these culture supernatants and analyzed and compared. Although the EV concentration had increased strongly, their ADAM10 content, typically found in melanoma cell-derived EV (9, 16), was reduced and correlated inversely with the increasing ratio of co-cultured Huh7 cells (**Fig 1F**, lane 3, 4). Conversely, ADAM10 was strongly present in EV from non-co-cultured MCTC (**Fig 1F**, lane 2). To confirm this suppressive effect, we analyzed the content of <u>c</u>ytokines, <u>c</u>hemokines and soluble <u>f</u>actors (CCF) in EV from MCTC before and after co-culture by protein microarray. While EV derived from MCTC contained a rich CCF profile, this profile was suppressed when MCTC were co-cultured with Huh7 cells in a cell ratio-dependent manner (**Fig 1H**, grey boxes; original array in **Fig S1B**). On the other hand new factors were secreted upon co-culture. These additional factors (yellow boxes), which included IFNγ, could have been secreted by both cell lines. Clearly, however, these factors were produced as a consequence of the interaction of both cell types.

When we analyzed the miRNA content in EV secreted by co-cultured cells, we found an increase of MDM2/4 regulating and tumor cell suppressive miR-34a, miR-192 and miR-194, similar as in our recent study (9), which were present in only low concentrations in EV secreted by either cell line alone. Conversely miR-215 was not elevated. While we could not determine the cellular origin of those miRNAs, it seemed likely that they were produced by the Huh7 hepatocytes. Taken together, hepatocytes exerted a strong suppressive effect on MCTC, potentially executed at least in part by EV and associated factors.

#### Hepatocyte/ tumor cell co-cultures secrete miRNAs also found in tumor patients

To substantiate the assumption of liver-derived tumor-suppressive pEV, we wanted to expand our analysis on the interaction of tumor cells with cells of the innate immune system. We speculated that these co-cultures would (1), produce unique EV-associated factor profiles that should be detected in patient’s pEV and (2), would allow a deduction of its potential cellular origin if a similar interaction occurred in vivo. We systematically assessed EV miRNAs and CCF profiles from co-cultures (for 48h) of 3 different liver cell types (as in **Fig. 1B**) with 3 different primary melanoma lines, each in two different co-culture ratios (1:1, 2:1). The same analysis was performed using resting PBMC from two different donors at a higher ratio of 30:1, since some PBMC sub-fractions, e.g. dendritic cells, are low in number. The results were compared with factors found in pEV from 12 melanoma patients and 2 healthy controls. Melanoma patients were randomly selected from three clinical stage groups, namely **Re**-(patients with a <u>re</u>lapsing tumor), **HR**-(patients with a <u>h</u>igh risk for tumor relapse) and **LR**-patients (patients with a low risk for tumor relapse)). In Re-patients, a tumor relapse was detected upon routine clinical presentation (every 3 month), irrespectively of the original clinical stage. A summary of this analysis scheme is presented in **Fig. 2A**.

**Figure 2:**
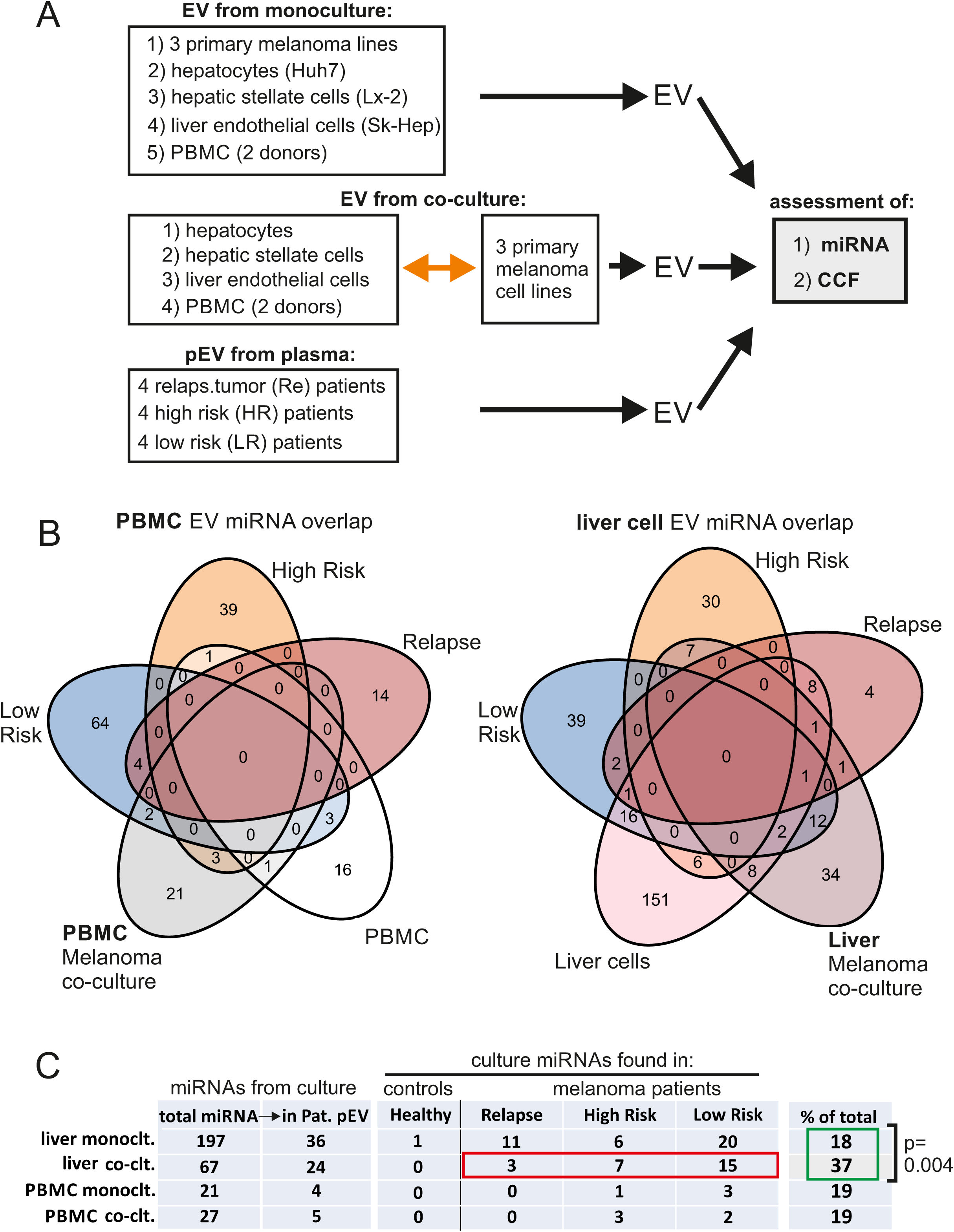
pEV-associated miRNAs secreted from liver/melanoma co-cultures are found in melanoma patients. **A** Schematic overview of the experimental procedure described in the text. EV were purified by differential centrifugation either from cell monocultures or co-cultures as indicated. Similarly pEV were purified from 12 melanoma patients in the 3 different clinical stages (see text for details). Micro-RNAs and protein content was extracted from the vesicles and analyzed by commercially available test systems (NanoString, protein arrays). **B** Venn diagrams depicting the presence of upregulated miRNAs in each patient group and cell culture, and their respective overlap. An up-regulation was defined as at least a two-fold increase over reference or category groups as explained in **Table S1**. The overlap of miRNAs between selected categories was inspected using Venn diagrams. **C** Summary of the Venn diagrams depicted in (B). Imbalances in the miRNA distribution were checked on overlap-derived contingency tables with the chi-squared test. This revealed that miRNAs found in liver co-cultures (a total of 67) are also found in pEV of melanoma patients, particularly HR and LR patients (red box). Compared to the presence of miRNAs from liver monocultures (a total of 197), this was significant (yellow box).

For miRNA analysis the NanoString technology was used, performed by a qualified operator (service facility). From these data sets, obtained from 14 individuals and 32 mono- and co-cultures, only miRNAs were included showing at least a twofold increase over their respective reference categories (as defined in **Table S1**). This procedure lead to a representative miRNA population for each patient and cell culture group as indicated by the Venn diagrams in **Fig 2B** (list of miRNAs in **Table S2**).

Analyzing the overlap between patient’s pEV and culture EV, we found that EV miRNAs from liver-melanoma co-cultures were significantly more present in pEV obtained from melanoma patients (in % of total) as compared to miRNAs from PBMC-melanoma co-cultures, or liver and PBMC monocultures (**Fig 2C**, green box). Furthermore, these miRNAs were not found in healthy individuals. For example, all liver co-cultures produced a total of 67 miRNAs (**Fig 2C**), which were at least 2-fold higher than in reference categories. Of those, 24 (37%) were found in patient’s pEV, namely 3 in Re-, 7 in HR- and 15 in LR patients (**Fig 2C**, red box). Thus, LR- and HR patients, who suppress tumor relapse, harbored considerably more of these miRNAs as compared to Re-patients. MiRNA from all other cultures were not found in higher numbers. These results supported the assumption that an interaction between liver cells and tumor cells occurred in tumor patients and induced the secretion of liver-derived pEV.

#### Tumor cell CCF secretion is suppressed following liver cell co-culture

Next we analyzed the CCF content of EV purified from the same in vitro cultures described in **Fig 2A**. While EV secreted from liver cell mono-cultures contained very few factors, EV from all three melanoma cell lines, and the above described MCTC line, contained a prominent and overall similar CCF profile. Notably, ICAM1, MIF and MCP-1 were secreted by all 4 lines (**Fig S2**).

Upon co-culture with liver cells these EV profiles changed significantly. Many of the factors secreted by tumor lines were strongly suppressed, mostly in a liver cell-ratio dependent manner as shown in **Fig 3A** for Huh7 hepatocytes (red boxes; quantification see in **Fig S3A**). Conversely, factors not detected in either monoculture were upregulated (blue boxes), again in a cell ratio-dependent manner. Analyzing the effects of all liver lines (**Fig S4;** quantification in **Fig S3B**), each cell line showed an individual pattern of suppression and secretion when co-cultured with each of the 3 melanoma lines (**Fig 3B**) and each pattern was similar against the different tumor lines. Together these data suggested that liver cells reacted actively and strongly to the encounter of tumor cells.

**Figure 3:**
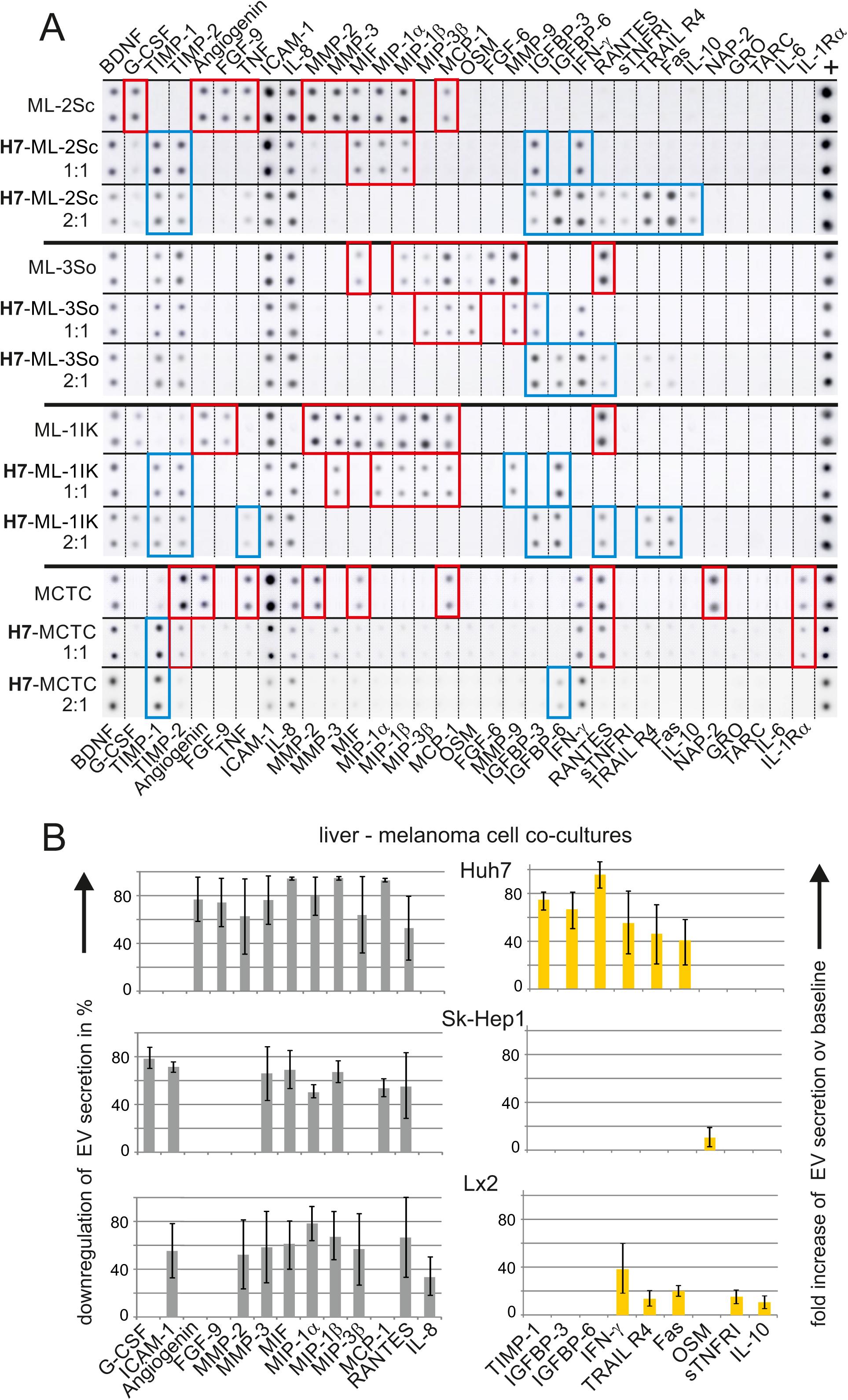
CCF factors detected in EV of liver/melanoma co-cultures. **A** Protein arrays of EV from melanoma cells before and after co-culture with Huh7 hepatocytes. Shown are original signals serving as the basis to calculate the pixel density and relative concentration of CCF in the indicated liver cell mono-cultures and liver/melanoma co-cultures. The cells were cultured or co-cultured for 48 h at a ratio of 1:1 and 2:1, before being purified and analyzed as described in **Fig S2**. The original signals were rearranged for better overview and comparison. Down-regulated melanoma-derived factors are indicated by red boxes, whereas up-regulated factors are indicated by blue boxes. **B** Summary of EV CCF regulation in liver/melanoma co-cultures. Based on the data/numbers shown in (A), mean values and error bars (student t-test) were calculated for the effect of each liver line on all 3 melanoma lines (MCTC not included). Only factors are shown and analyzed that were up or down regulated in any of the co-cultures. The gray bar diagrams depict the relative down regulation of EV secretion with respect to the levels seen in melanoma mono-cultures. The yellow bars depict the relative up-regulation of new CCF factors, expressed as fold increase over baseline levels in melanoma, liver and PBMC mono-cultures.

#### Tumor cell CCF secretion is suppressed following PBMC co-culture

A similar observation was made with PBMC. Resting PBMC did not secrete EV with a notable CCF content (**Fig 4A**, upper graphs). Again, this changed significantly when tumor cells were co-cultured. Some of the tumor-derived factors were completely suppressed (**Fig 4A**, lower graphs, red boxes), while other factors not present in PBMC or tumor cell monocultures were strongly upregulated (blue graphs). Both PBMC donors reacted similarly to all co-cultured tumor lines (**Fig 4B;** quantification in **Fig S5**). The strong PBMC reaction was in part due to the HLA mismatch of the interacting tumor cells. However, since tumor cells are immunologically recognized by the host immune system, the here measured pEV secretion may represent the maximal response of a tumor cell/ PBMC encounter.

**Figure 4:**
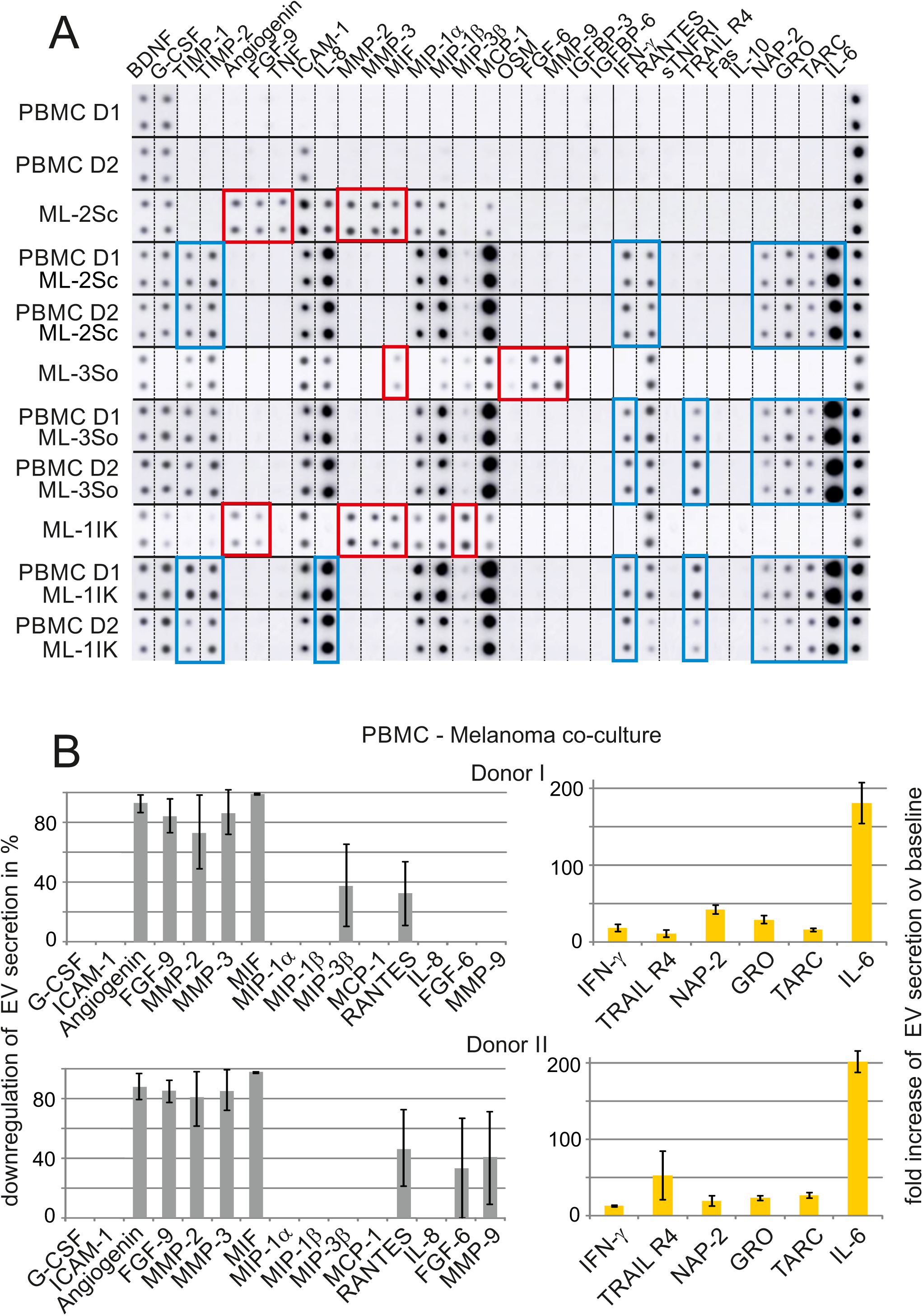
CCF factors detected in EV from PBMC/melanoma co-cultures. **A** Protein arrays of EV from melanoma cells before and after co-culture with PBMC. Shown are rearranged original signals as in **Fig 3A** of CCF protein arrays with purified EV from PBMC mono- and PBMC/melanoma co-cultures. Only resting PBMC were used. The cells were cultured or co-cultured for 48 h at a ratio of 30:1, before being processed as in (A). Down-regulated melanoma-derived factors are indicated by red boxes, whereas up-regulated factors are indicated by blue boxes. **B** Summary of EV CCF regulation in PBMC/melanoma co-cultures. Based on the data/numbers shown in (A), mean values and error bars (student t-test) were calculated for each factor secreted by both PBMC donors upon co-culture with all 3 melanoma lines. Only factors are shown and analyzed that were up or down regulated in both co-cultures. The gray bar diagrams depict the relative down-regulation of EV CCF secretion with respect to the levels seen in melanoma mono-cultures. The yellow bars depict the relative up-regulation of new CCF factors in co-cultures, expressed as fold increase over baseline levels in melanoma and PBMC monocultures.

#### CCF comparison from pEV and cell culture EV suggests tumor- and host-derived factors

The presence of EV CCF factors in the various in vitro cultures was summarized and compared with the relative presence of factors detected in patient’s pEV. For this the average level for each factor in each patient group was determined and color coded (**Fig 5A** left rows; primary data of patients pEV in **Fig S6**; normalized individual patient’s results in **Fig 5B)**. Deducing from this synopsis, factors were deemed “tumor-associated” when they were (1) present in Re- and absent in LR patients, (2) were produced by tumor cells and not by liver or PBMC monocultures, and (3) when they were absent and/or suppressed in liver and/or PBMC co-cultures. For example, MMP9 was strongly present in 4/4 Re-patients, but almost absent in HR (1/4) and LR patients (0/4). MMP9 was not produced by the liver or PBMC monocultures, but produced by all tumor lines. MMP9 was not present in liver or PBMC co-cultures, as it’s secretion from tumor cells was likely suppressed. Conversely, factors were deemed “liver/PBMC-associated”, when they were (1) not detected in tumor cell monocultures, but (2) were present and (3) not suppressed in liver/PBMC co-cultures. For example, IFNγ was not produced in tumor monocultures, but strongly present and not suppressed in the liver/PBMC - tumor co-cultures. Hence, IFNγ likely derived from liver cells and/or PBMC. Using this algorithm, we assumed/assigned 6 of 24 factors to derive from tumor cell activity and 5 factors to derive from PBMC and/or liver cell activity (**Figure 5A**, right column).

**Figure 5:**
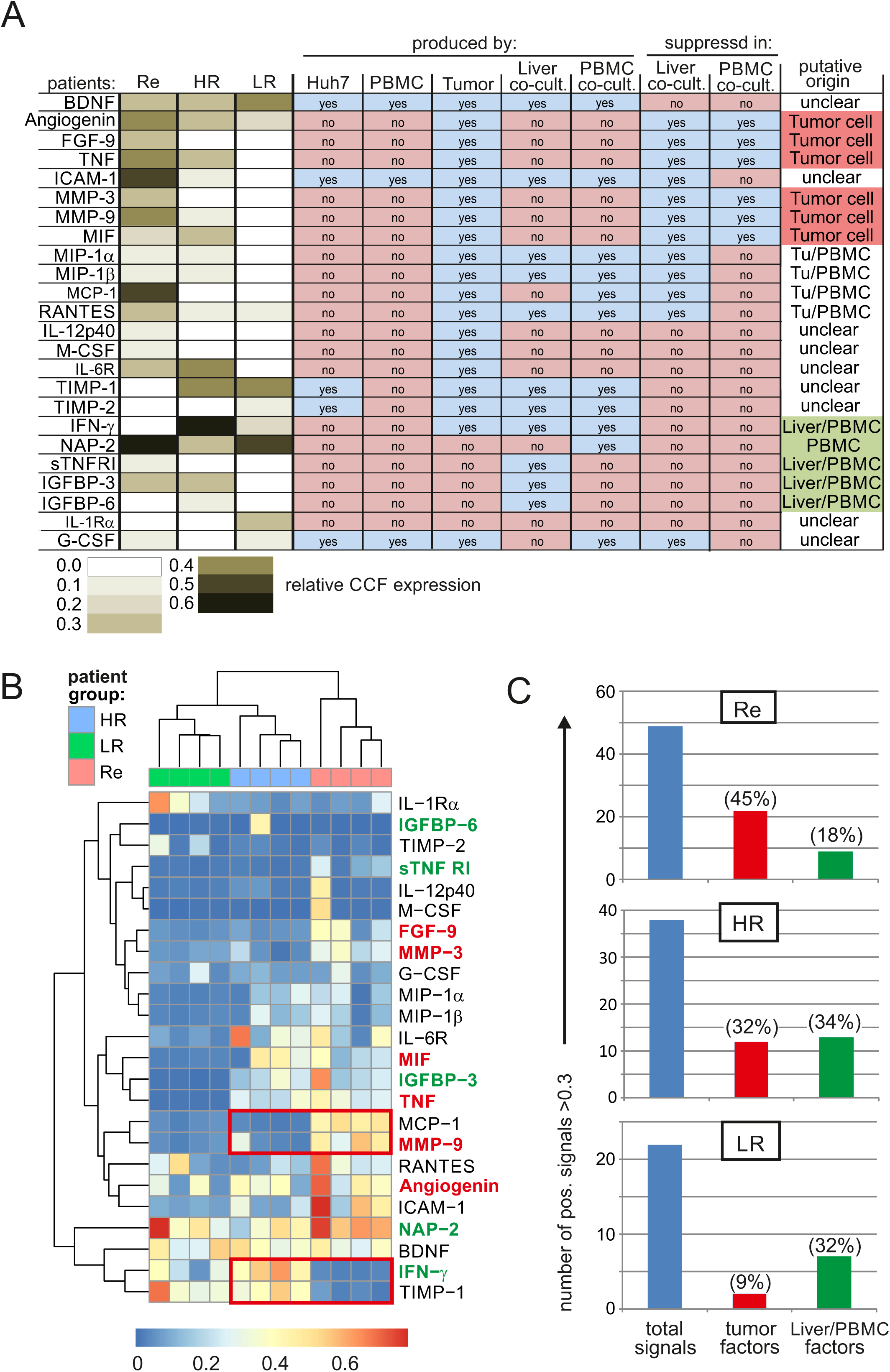
Identification of tumor- and liver/PBMC-derived factors in pEV of melanoma patients correlating with the clinical stage. **A** pEV factors in melanoma patients and their presence/absence in cell culture derived EV. On the left factors are depicted found in patient’s pEV. Their relative/average levels were calculated on the basis of protein array signals (**Fig S6**) using ImageJ. Values were normalized with respect to the internal positive controls and different levels of expression were color coded for each patient group. The presence/absence (yes/no) of these factors in all cultures and co-cultures was determined accordingly, based on results in **Fig 3** and **4**. Based on this information their putative cellular origin was deduced (right row), in an algorithm explained in the text. **B** The number of CCF factors detected in melanoma patients correlates with their clinical stage. The CCF factors quantified in patient’s pEV were depicted by heat map. Factors depicted in red letters are of putative tumor cell origin and factors in green letters of putative liver/PBMC origin as predicted by the algorithm in (A). Red boxes are explained in the text. **C** The presence/absence of pEV factors in melanoma patients pEV correlates with their clinical stage and tumor relapse. The blue bars depict the number of all positive CCF signals found in each melanoma patient group (4 patients in each group). The red and green bars depicts the number of tumor-associated factors (red) and liver/PBMC associated factors (green) in each patient group as determined in (A). The respective percentages were calculated with respect to the total number of signals and depicted in parenthesis.

#### CCF factors in pEV deemed tumor- or host-derived correlate with clinical stage

To validate the assignment of these factors, we asked whether they correlated with the clinical stage (**Fig 5B** and **C**). First we noticed that the presence, respectively absence, of two of the assigned markers (MMP9 and IFNγ) correlated with tumor relapse while their inverse presence/absence in HR and LR-patients correlated with tumor control. The same was true for two non-assigned factors, TIMP-1 and MCP-1 (**Fig 5B**, red boxes).

We then counted all positive signals (>0.3 relative intensity in **Fig 5B**) in each patient group (**Fig 5C**, blue bars), as well as the number of tumor- and liver/PBMC-associated signals (**Fig 5C** red and green bars). First, all Re-patients harbored significantly more factors in their pEV (49) than HR (37) and LR-patients (22). Second, in Re-patients a surprisingly high number of these signals represented tumor-assigned factors (21/49: 45%), but only a low number were liver/PBMC-associated (9/45:18%). In HR patients the ratio of tumor- and liver/PBMC-associated factors was evenly distributed (32% vs 34% respectively), whereas this ratio was inverted in LR patients (9% vs. 32%). In summary, our assignment of factors correlated surprisingly well with the clinical stage and risk for tumor relapse.

## Discussion

We have recently demonstrated that post-operation tumor patients harbor elevated levels of tumor cell suppressive pEV. Based on indirect evidence, we now conclude that liver cells, beside PBMC, are a potential source of these cancer-induced pEV. Surprisingly, we also found evidence for a strong presence of tumor-derived pEV, implying that residual melanoma cells are potentially more numerous and more active than anticipated. Finally, we present preliminary results from patient material, indicating that patterns of pEV-derived factors could serve to predict cancer relapse.

At present no technology exists that could reliably trace pEV to their cellular origin, since none of the proteins and factors contained therein are truly unique for a specific organ or tissue. Hence, our assignment of factors to tumor- or liver/PBMC cell origin on the basis of their respective absence/presence in pEV and EV from in vitro cultures (**Fig 5A**) remains an assumption. Nevertheless, there are several arguments that support our approach and conclusion/hypothesis. First, liver cells and PBMC reacted strongly upon encounter of tumor cells as demonstrated here (e.g. **Fig 1E**). Even when we take the HLA mismatch in our system into account, a reduced but similar reaction is likely to occur in vivo. Second, this reaction did not kill tumor cells, as would have been expected for a nonspecific interaction, but resulted in a strong and seemingly specific tumor cell-suppressive activity including the upregulation of p53/β-catenin suppressive miRNAs (9) (**Fig 1F**-**H**). Third, we analyzed patterns of factors and micro-RNAs and not only single markers (**Fig 5B**). Finally, the presence/absence of tumor- and liver/PBMC-associated pEV factor patterns, including miRNAs, correlated surprisingly well with the clinical stage and tumor relapse (**Fig 2C, 5B and C**).

Growing tumors may shed large amounts of cells into circulation (17). Not all of these cells will be readily recognized and killed by the adaptive immune system. Hence innate immune cells are likely the first line of defense against a growing and immunologically changing tumor (18). It is plausible that the liver plays an important role in this scenario as it is a large immune sensing organ screening the whole blood volume every 3 minutes (19). In addition, it has the size and secretory capacity to supply pEV in high numbers. Liver cells may sense circulating tumor cells through direct contact, as demonstrated here in vitro, or by secreted factors, e.g. cytokines and tumor vesicles, and potentially through tumor cell typical secretion of RNA elements and endogenous retroviruses (20, 21). Hence, remaining tumor cells after R0 surgery (CTC/DTC) may stimulate liver cells and PBMC for, in general, lower but persistently elevated pEV levels.

The strong presence of markers, particularly in relapsing (Re) patients (**Fig 5A**), that we deemed as being secreted from tumor cells, was surprising, because the number of CTC/DTC is generally considered to be very low (22). It is assumed that most CTC/DTC die rapidly, as shown in animal models (23, 24). However, the situation in HR- and particularly in relapsing patients may be somewhat different from experimental systems. CTC/DTC cell clones may have learned to proliferate while escaping immune surveillance. In addition, the growing tumor mass in Re-patients likely contributes significantly to the presence of tumor cell-derived pEV. In any scenario, such pEV secreting tumor cells, and potentially also their tumor microenvironment (25, 26), would have to proliferate to significant numbers in order to produce the amount of pEV-associated factors measured in this study in plasma.

The pattern of tumor and immune-derived factors secreted by pEV may provide a snapshot-like insights into the battle between the immune system and tumor cells. Our preliminary analysis of these patterns in three different clinical stages of melanoma supported this notion and revealed an astonishing discriminatory power of these patterns (**Fig 5B**). This was particularly apparent between high-risk patients that controlled their CTC/DTC tumor compartments and those patients experiencing a tumor relapse. The presence/absence of four factors (MCP-1, MMP-9, IFNγ and TIMP-1) correlated stringently with tumor control and tumor relapse. With the help of artificial intelligence and higher case numbers more information is likely to be extracted from these patterns than demonstrated here. Hence, such pEV factor patterns could become import diagnostics tools in the management of cancer patients.

In summary our results imply that tumor cells induce a strong reaction of the innate immune system, including liver cells. Notably, our results also imply that CTC/DTC are more prevalent and/or more active than currently believed, otherwise we would likely not see the rich CCF and miRNA response in patient’s pEV. Finally, the profile of pEV-associated factors may help to estimate the relative risk for melanoma relapse. However this requires the analysis of large numbers of patient plasma samples.

## Material and methods

### Cell lines and primary cells

#### Cell lines

Liver cell lines Huh7 and Sk-Hep-1 (kindly provided by P. Knolle, Technische Universität München) were grown in DMEM (Sigma-Aldrich) supplemented with 10% Fetal calf serum (FCS, Sigma-Aldrich) and 1% penicillin-streptomycin (Lonza). Sk-Hep1 cells were additionally maintained in 40 M-mercaptoethanol (Carl Roth). LX-2 cells were provided by SL. Friedman (Icahn School of Medicine, New York) and cultured in DMEM high glucose (Life Technologies) supplemented with 2% FCS, 1% penicillin-streptomycin. All cells were grown at 37°C under 5% CO_2_. **Peripheral blood mononuclear cell (PBMC) preparation**: Leukoreduction system chambers (LRSCs) from healthy donors were acquired after plateletpheresis. The resulting platelet free cell sample was diluted 1:2 in PBS and the PBMC containing buffy coat was isolated after density gradient centrifugation on Lymphoprep (Axix Shield 1114544). **Generation of immature/mature Dendritic cells (DC)**: Monocytes were isolated from PBMCs using BD IMag Anti-Human CD14 Magnetic Particles (BD Biosciences 557769). 6.0 × 10^6^ monocytes were seeded in a 6 well plate in RPMI supplemented with 1% human serum (Sigma-Aldrich). Monocyte-derived DC were generated adding 800 IU/ml of recombinant GM-CSF and 250 IU/ml of recombinant IL-4 (both from CellGenix). For EV isolation (see below) immature DC were washed and 24 h later the supernatant was harvested (10 ml). To generate mature DC, immature DC cultures were supplemented for 24 h with LPS (100 ng/ml) or a maturation cocktail (200 IU/ml IL-1ß, 1,000 IU/ml IL-6 (both from CellGenix), 10 ng/ml TNF (beromun; Boehringer Ingelheim) and 1 µg/ml Prostin E2 (PGE2, Pfizer). Subsequently cells were washed and EV supernatants (10 ml) were collected 24 h later for EV isolation. **Generation of Macrophages:** Monocytes were separated from the non-adherent fraction (NAF) by plastic adherence on cell culture flasks and cultured in RPMI supplemented with 1% human serum and 1% of penicillin/streptomycin. On days 1, 3, 5, 7 and 9 medium was supplemented with 800 IU/ml of GM-CSF. On day 11, medium was removed, cells were washed and 20 ml of RPMI supplemented with 1% of EV depleted human serum was added. After 24 h the supernatant was harvested and EV were isolated. For all procedures see also (10). **MCTC cell line**: From 30 ml blood of a melanoma patient the CD45-positive cells were depleted using CD45 RosetteSep (Stemcell Technologies) according to manufacturer’s instructions. The remaining cells were stained with MCSP-APC and MCAM-FITC antibodies (both from Miltenyi) and DAPI (Thermo Fisher) for dead cell exclusion. MCSP-positive and/or MCAM-positive cells were then sorted on a FACS Aria SORP (BD) cell sorter and seeded in RPMI cell culture medium with 20% human pooled serum. Medium was replaced on a regular basis and cells showed first signs of growth after several weeks. At the time the CTC cells were obtained the patient was tumor free. **Melanoma lines ML-1IK, ML-2Sc and ML-3So**: Melanoma tumor cell lines were generated as described before (16). Briefly, fresh tumor biopsies were obtained directly after surgery. A single cell suspension was produced by mechanical dissociation and enzymatic digestion with DNAse and collagenase. Cells were seeded in RPMI supplemented with 20% human serum into 6-well plates. Passaging of cells was performed according to cell density.

### EV depletion of FCS and human serum for cell culture

To assure that EV generated from cell culture were not contaminated by outside sources, heat-inactivated FCS and human serum for medium supplementation were depleted of bovine EV by ultracentrifugation for 18 h at 110,000 g and 4 °C before use.

### Antibodies and Reagents

Primary antibodies were used at 1–2 μg ml^−1^ for immunoblotting: anti-ADAM10 (mouse monoclonal, Abcam ab73402), anti-CD63 (mouse monoclonal, BD Biosciences 556019), anti-CD81 (mouse monoclonal, BD Biosciences 555675), anti-Haptoglobin (rabbit polyclonal, Biozol, GTX 112962-25). The following secondary antibodies were used: Alexa Fluor 488 goat anti-mouse and Alexa Fluor 555 goat anti-rabbit IgG (both from Life Technologies) and anti-mouse IgG-HRP conjugate and anti-rabbit IgG-HRP conjugate (both from Cell Signaling).

### Patient material

#### Cohort 1 (Fig 1A)

This is the same patient cohort described in Lee et al., 2019. Briefly, Plasma samples were obtained from patients attending the outpatients departments at the University Hospital Erlangen after signing an informed consent. The study was approved by the local ethics committee in Erlangen (Nr. 4602). Patients were assigned to the respective study groups based on their clinical stage (27). R0 operated patients were subdivided into high risk (HR) (stage II-IV) and low risk (LR) patients (stage I). T patients harbored tumor metastases (clinical stage III and IV), or primary tumors (clinical stage I – II) before surgery.

#### Cohort 2 (Fig 2, 4, 5)

Plasma samples were obtained as for cohort 1 and patients, with the exception of Re-patients, were categorized in LR and HR groups as described for cohort 1. In Re patients (relapsing patients), the blood sample was taken upon tumor relapse, detected at routine clinical presentations (every 3 month) or upon ad hoc presentations after patients detected a new growing lump. A summary of the patient data is listed in **Table S3**.

#### CAT plots (patient cohort 1)

The CAT (correspondence at the top) plots were generated with miRNA data obtained from patient cohort 1 and primary immune cells (mature and immature dendritic cells, macrophages). The miRNA assessment by a commercial provider was described in (9). The miRNA extraction is described below. For the CAT plot analysis we adapted a method used previously to compare the agreement in measurements between microarray platforms (28). Each trace in the graphs (**Fig 1A**) corresponds to one comparison between two groups, as indicated by the legend. Specifically, the traces show the percentage of common miRNAs in two group rankings plotted against rank. To rank the miRNAs in each group, the mean (m) and standard deviation (SD) of miRNA EV concentrations were estimated as follows: For all groups (cells) there was only one measurement available (n=1), so m was set to signal intensity and SD to signal error, as recorded by the instrument. A score S for every microRNA was calculated according to S = m / SD, and miRNAs were ranked by sorting S in descending order for each group.

### miRNA assessment and analysis for patient cohort 2

The miRNA content of pEV (see below), was reverse transcribed into cDNA as described previously (9). Subsequently the miRNA quantification was performed on the NanoString platform according to the manufacturer’s instructions using the respective miRNA assessment and quantification kit (GXA-MIR3-12). Resulting counts were subjected to background correction based on negative controls and global normalization in R (R Core Team (2018). R: A language and environment for statistical computing. R Foundation for Statistical Computing, Vienna, Austria. https://www.R-project.org/) with the package NanoStringQCPro (https://doi.org/doi:10.18129/B9.bioc.NanoStringQCPro>). An additional round of quantile normalization was applied separately to the patient samples (n = 20) and the cell population samples (n = 44), respectively. Afterwards, replicate means were calculated for each patient and culture category and separated into three reference groups: patient-derived blood plasma, in vitro monocultures and in vitro co-cultures. In each of these groups independently, miRNAs were analyzed for overrepresentation by checking if the normalized expression value exceeded the highest value from the reference categories (**Table S1**) by at least two-fold. MiRNAs for which this test turned out positive were deemed to be associated with the corresponding category. Using R and the package VennDiagram (https://CRAN.R-project.org/package=VennDiagram), the overlap of associated miRNAs between selected categories was then inspected with Venn diagrams (**Fig 2B**), and imbalances in the distribution were checked on overlap-derived contingency tables with the chi-squared test.

### Isolation and purification of EV and pEV

EV and pEV purification was performed essentially as described previously (10). Briefly, cell culture supernatants were collected after 48 h and centrifuged for 20 min at 2,000 g, 30 min at 10,000 g and ultra-centrifuged for 1 h at 100,000 g. Pellets were resuspended in 35 ml PBS and centrifuged at 100,000 g for 1 h. Pellets were resuspended in 100 μl PBS and considered as EV preparations. For pEV purification, 10 ml blood plasma was diluted with 10 ml PBS and centrifuged for 30 min at 2,000 g, 45 min at 12,000 g and ultra-centrifuged for 2 h at 110,000 g. Pellets were resuspended in 10 ml PBS and centrifuged at 110,000 g for 1 h. Pellets were again resuspended in 100 μl PBS and considered as EV preparations. These Pellets were solubilized in SDS sample buffer or re-suspended in 100 μl PBS and aliquots were analyzed by immunoblotting or Cytokine/Chemokine/soluble Factor (CCF) protein array (see below).

### Human cytokine/chemokine/soluble factor (CCF) array

Purified EV or pEV corresponding to equal volume of cellular supernatant (in general 60 ml from 10 mio. transfected cells) or equal plasma volume were applied to the RayBio Human Cytokine Array C-S (Hölzel Diagnostika, AAH-CYT-1000-2) according to the manufacturer’s instructions. A minimum of 20 μg EV proteins was used per filter incubation.

Spot signal intensities were quantified with the ImageJ plugin Protein Array Analyzer. For cross-blot normalization, subtraction of the average intensity of blank spots (n = 14) and division by the average of the positive controls was performed. A list of factors of interest was then selected and plotted in a Euclidean-distance heat map using the library heatmap in R (https://CRAN.R-project.org/package=pheatmap)

### Quantitative PCR amplification

The procedure was described in detail in (9). Reverse transcription of extracted pEV RNA was performed using the commercially available QantiTect Reverse Transcription kit (Qiagen, Cat. No: 205311) or TaqMan® MicroRNA Reverse Transcription Kit (ThermoFisher, Cat. No: 4366596) using commercially available TaqMan® MicroRNA Assays (ThermoFisher, Cat. No: Cat. # 4427975). For amplification of miRNAs, qRT-PCR was performed using TaqMan® MicroRNA Assays (ThermoFisher, Cat. No: Cat. # 4427975) with a Rotor-Gene Probe PCR Kit (Qiagen, Cat. No: # 204374) according to the manufacturer’s instructions on a Qiagen Rotor-Gene Q real time PCR-cycler.

### Particle quantification

Sucrose-purified pEV were diluted 1:1,000 in PBS. The pEV numbers were quantified via particle tracking analysis on a commercially available ZetaView® particle tracker from ParticleMetrix (Meerbusch, Germany) using a 10 µl aliquot of the diluted samples. The concentration of pEV was calculated based on the dilution factors.

### Statistical analysis

Data were statistically evaluated using Student’s t-test or One-Way ANOVA subsequently followed by Tukey’s honest significant difference test when applicable.

### Data Deposition

The miRNA data sets were deposited at NCBI GEO. ID: GSE100508.

## Acknowledgement

This work was supported by the German Federal Ministry of Education and Research (BMBF) (01GU1107A) and MelEVIR (031L0073A) (JH.L.) and by the IZKF Erlangen (Interdisziplinäre Zentrum für Klinische Forschung). J.V. and M.E. were supported by the BMBF as part of the projects eBio:miRSys (0316175A) and eBio:MelEVIR (031L0073A). J.V-G. is also funded by the Elan Funds of the Medical Faculty of the Friedrich-Alexander-University of Erlangen-Nürnberg (FAU) and the DFG through the project SPP 1757/1 (VE 642/1-1).

## Author contributions

The project was designed and coordinated by A.S.B. Substantial contributions to the work were made by J-H.L. (pEV purification, transfections, Western blots, protein array and pEV analysis); M.E. and J. V. (Bioinformatics); K.B. (pEV purification).

## Figure Legends for Supplement Figures S1 - 6

**Figure S1:**
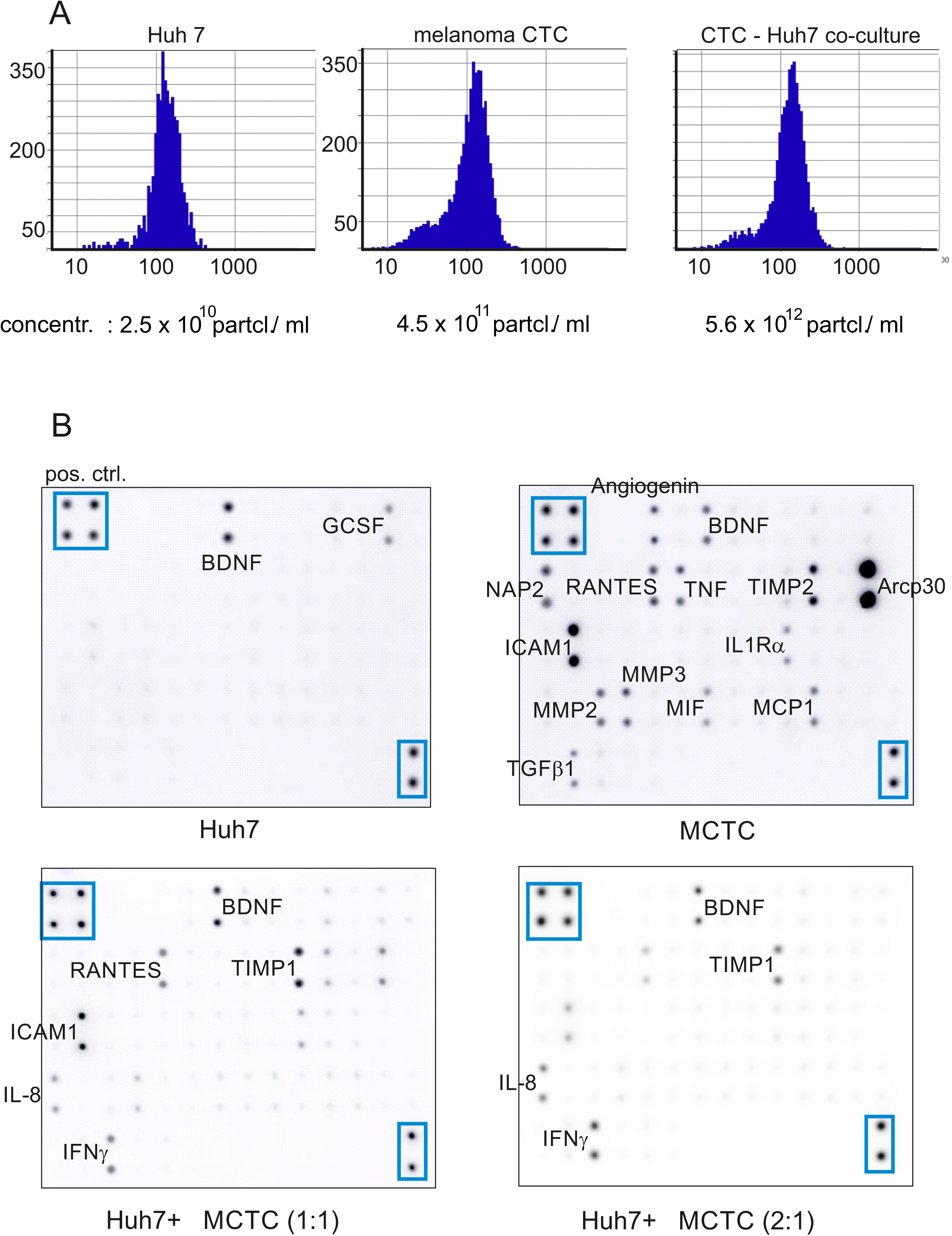
Co-culture of Huh7 and MCTC cells induces MCTC-suppressive EV secretion. **A** Particle/Vesicle number analysis derived from culture supernatants described in **Fig 1** co-culturing Huh7 hepatocytes with melanoma CTC cells (MCTC). Particle/Vesicle numbers were assessed by ZetaView® nanoparticle tracker. Shown is one representative experiment of two performed. **B** Original data (protein array by RayBiotec) serving to calculate the relative decrease of EV-associated <u>c</u>hemokines, <u>c</u>ytokines and soluble factors (**CCF**) after co-culture of Huh7 hepatocytes and CTC melanoma cells in **Fig 1G**. The cell lines were cultured in different ratios (1:1; 2:1).

**Figure S2:**
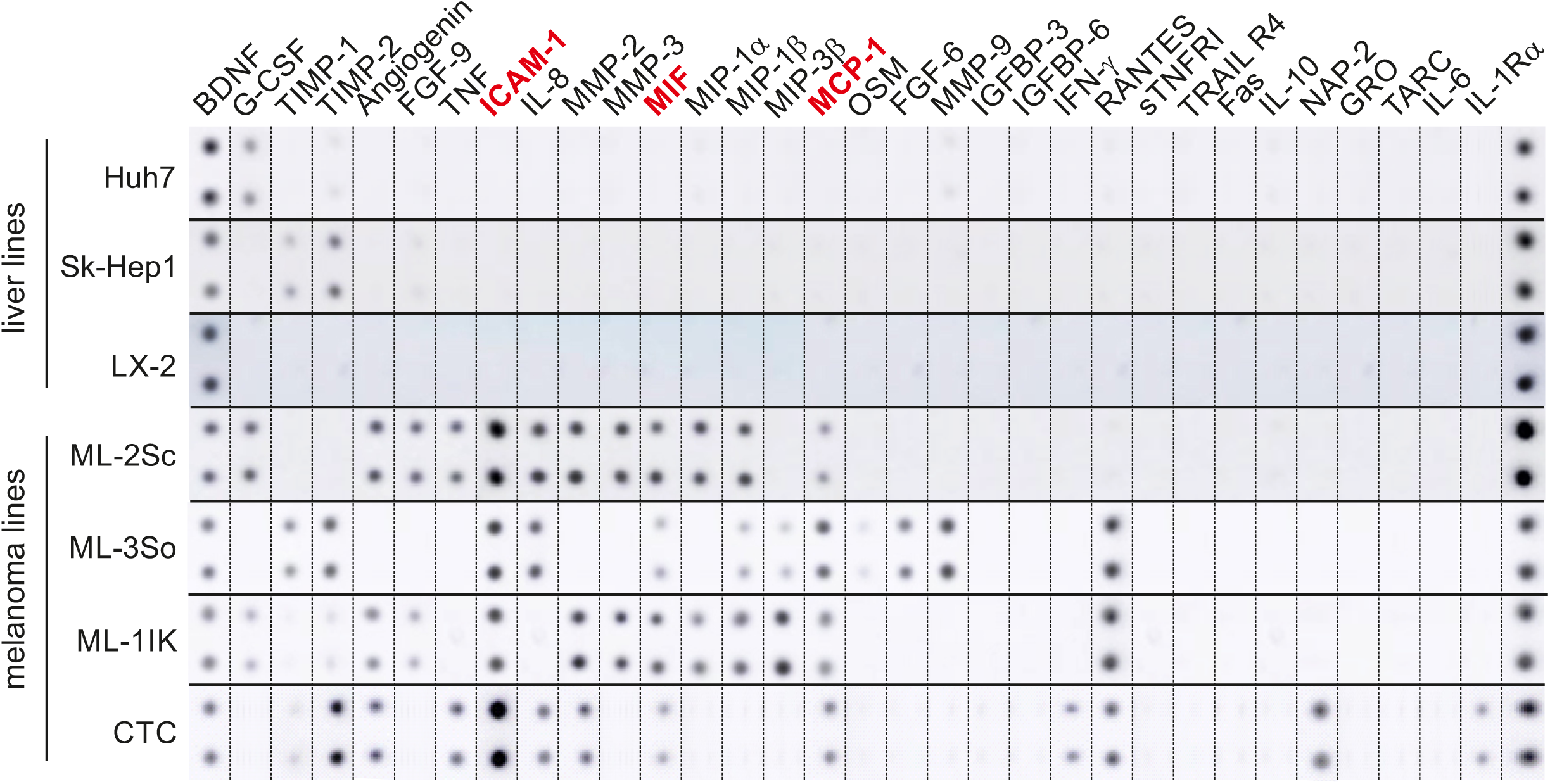
<u>C</u>hemokine<u>C</u>ytokine/soluble <u>F</u>actors (CCF) detected in EV from liver and melanoma mono-cultures. After 48h of cell culture of the indicated cell lines, supernatants were harvested and EV were purified by differential centrifugation. Subsequently all harvested EV were analyzed for the presence of CCF by a commercial array (Ray Biotech). Shown are the original signals which were rearranged for better overview and comparison. Factors depicted in red were detected in all 4 melanoma lines. The positive control (last row) is an internal control of the array confirming and measuring the reactivity of the secondary antibody, used to detect the specific signals. This internal control was used to calculate the relative pixel density of the signal.

**Figure S3:**
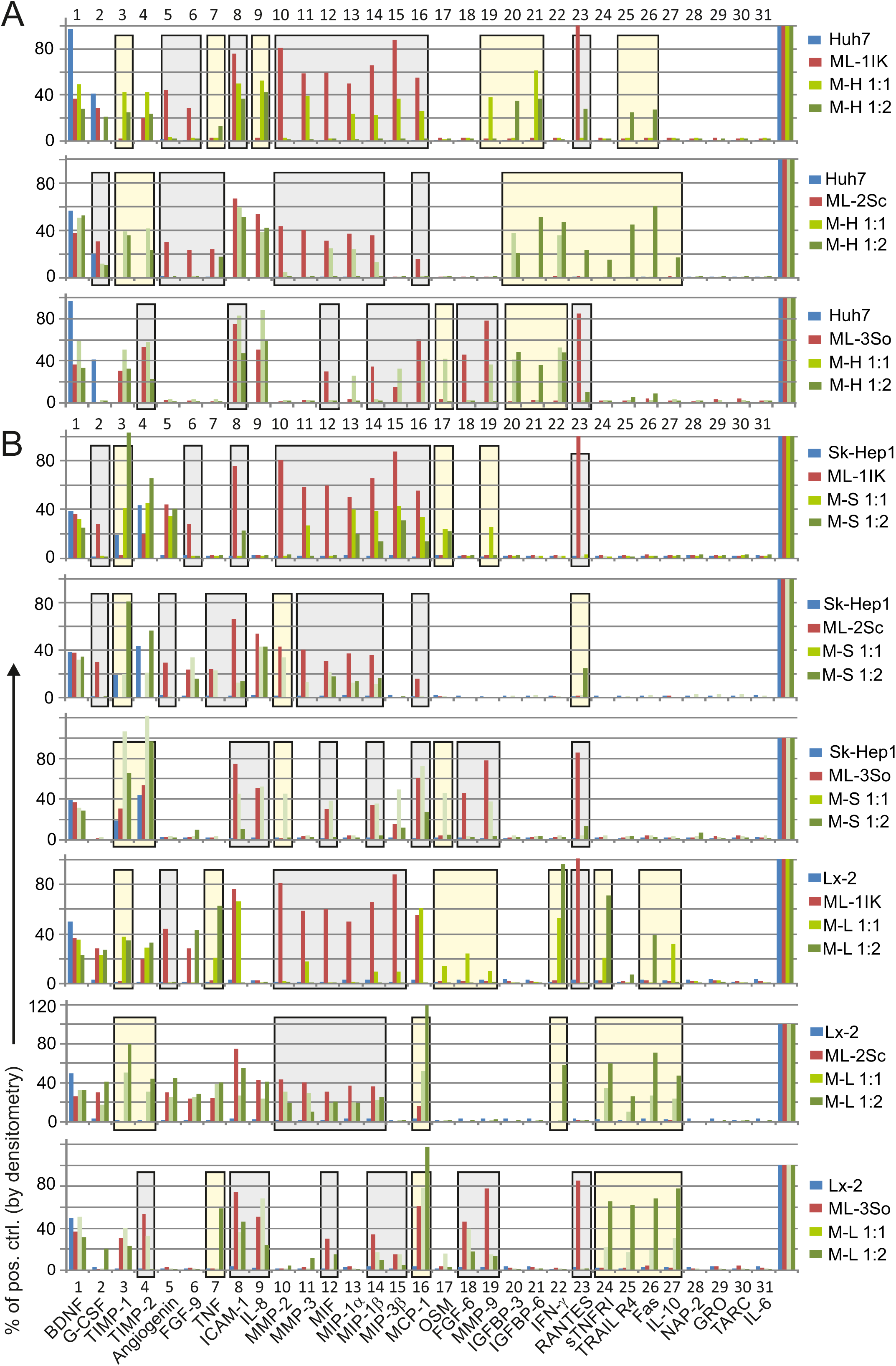
EV CCF levels in liver/melanoma co-cultures. **A** Quantification of CCF levels in Huh7 hepatocyte mono- and co-cultures with 3 melanoma cells. Original array signals of the EV CCF analyses (see **Fig 3A**) were quantified by densitometry and calculated in % from the positive controls (% of pos. ctrl.) for each blot. The positive control was set as 100 percent and is indicated by the 4 bars at the right end of the graphs. Only factors were analyzed that gave a positive signal in any of the in vitro cultures. Huh7 were co-cultured with 3 different melanoma cells (ML-1IK; ML-2Sc; ML-3So). Gray shaded areas indicate factors downregulated from concentrations recorded in melanoma monocultures (red bars) to levels seen in Huh7-melanoma co-cultures (green bars) with different cell number ratios (light green bars: 1:1, green bars: 2:1). Yellow shaded areas indicate factors that were not recorded in monocultures and significantly upregulated in co-cultures. On top of some graphs numbers were placed to better localize the position of individual CCF factors (see bottom of graphs). **B** Quantification of CCF levels in liver endothelial cells (Sk-Hep) and hepatic stellate cells (LX-2) with 3 melanoma cells. The analysis was performed as described for (A). The original array signals are shown in Fig S4.

**Figure S4:**
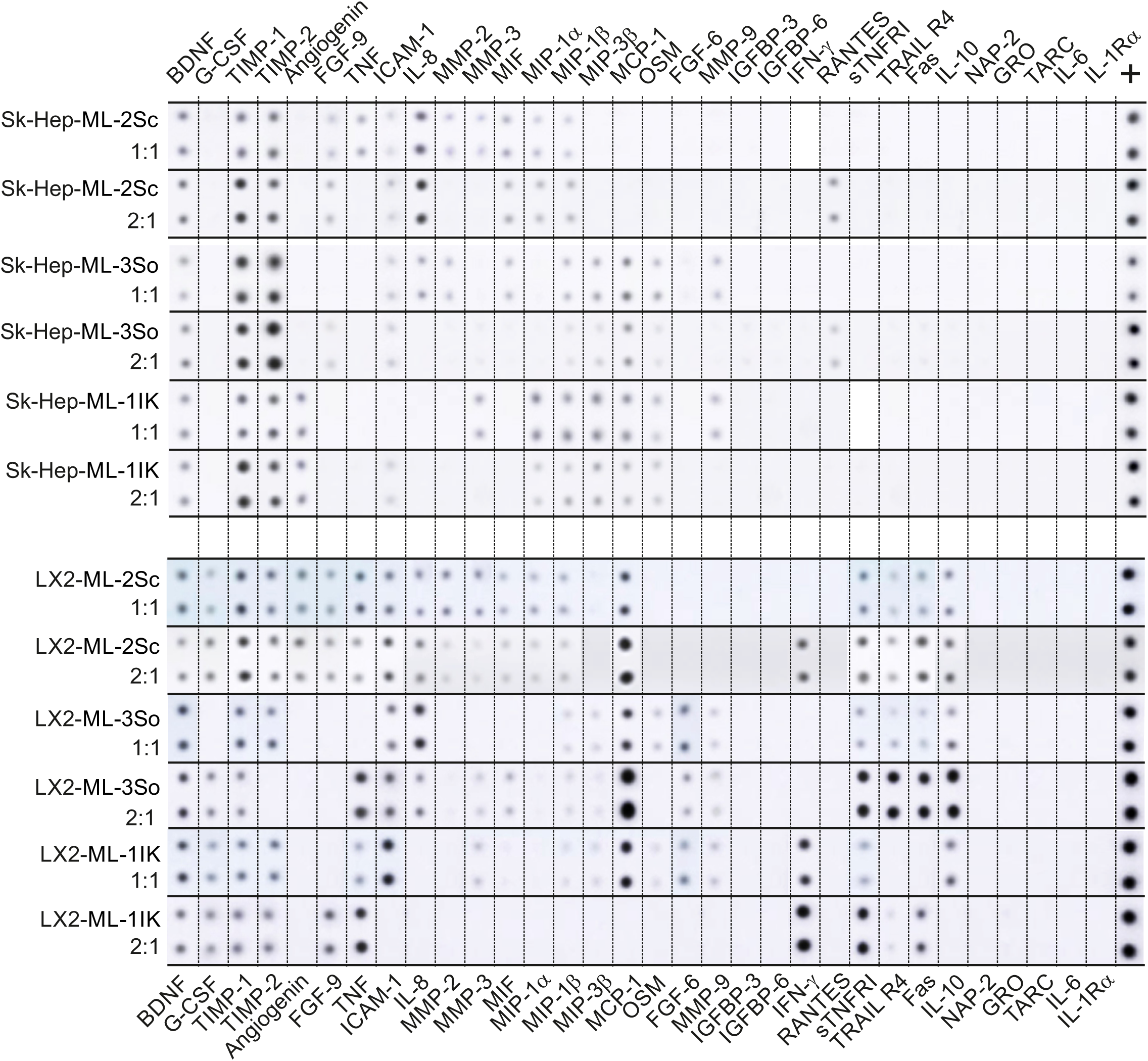
Original array signals of EV CCF analyses from liver-melanoma co-cultures Protein arrays of EV from melanoma cells before and after co-culture with SkHep and LX-2 liver cells. Shown are original signals serving as the basis to calculate the pixel density and relative concentration of CCF in the indicated liver cell mono-cultures and liver/melanoma co-cultures (**Fig S3)**. The cells were cultured or co-cultured for 48 h at a ratio of 1:1 and 2:1, before being purified and analyzed as described in **Fig S2**. The original signals were rearranged for better overview and comparison.

**Figure S5:**
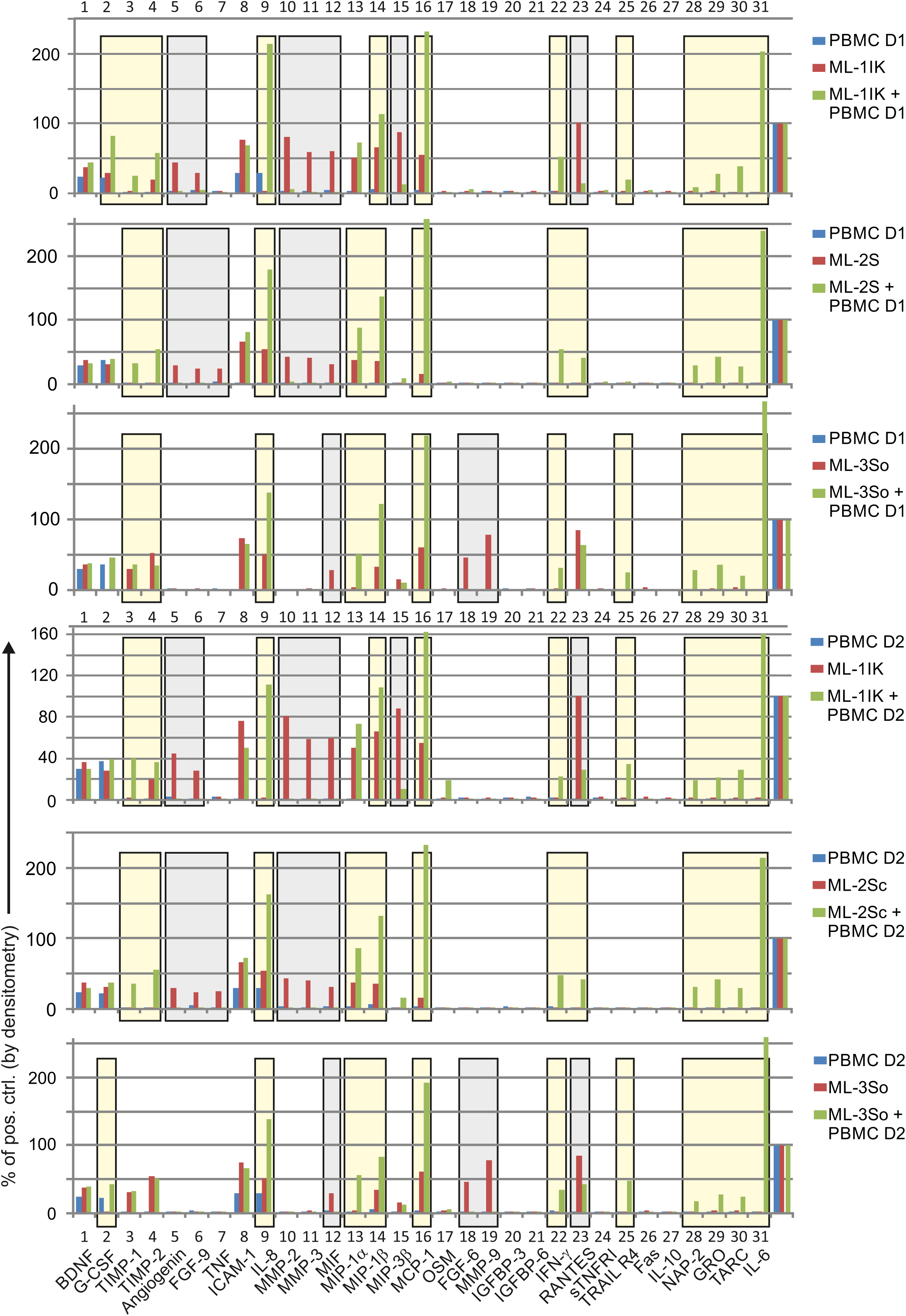
EV CCF levels in PBMC/melanoma co-cultures. Quantification of CCF levels in PBMC mono- and co-cultures with 3 melanoma cells. Original array signals of the EV CCF analyses (see **Fig 4A**) were quantified by densitometry and calculated in % from the positive controls (% of pos. ctrl.) for each blot. The cells were cultured or co-cultured for 48 h at a ratio of 30:1, before the supernatants were harvested and EV were purified by differential centrifugation and analyzed as described in **Fig EV2**. The positive control was set as 100 percent and is indicated by the 4 bars at the right end of the graphs. Only factors were analyzed that gave a positive signal in any of the in vitro cultures. PBMC from each of 2 donors were co-cultured with 3 different melanoma cells (ML-1IK; ML-2Sc; ML-3So). Gray shaded areas indicate factors downregulated from concentrations recorded in melanoma monocultures (red bars) to levels seen in PBMC-melanoma co-cultures (green bars). Yellow shaded areas indicate factors that were not recorded in monocultures and significantly upregulated in co-cultures.

**Figure S6:**
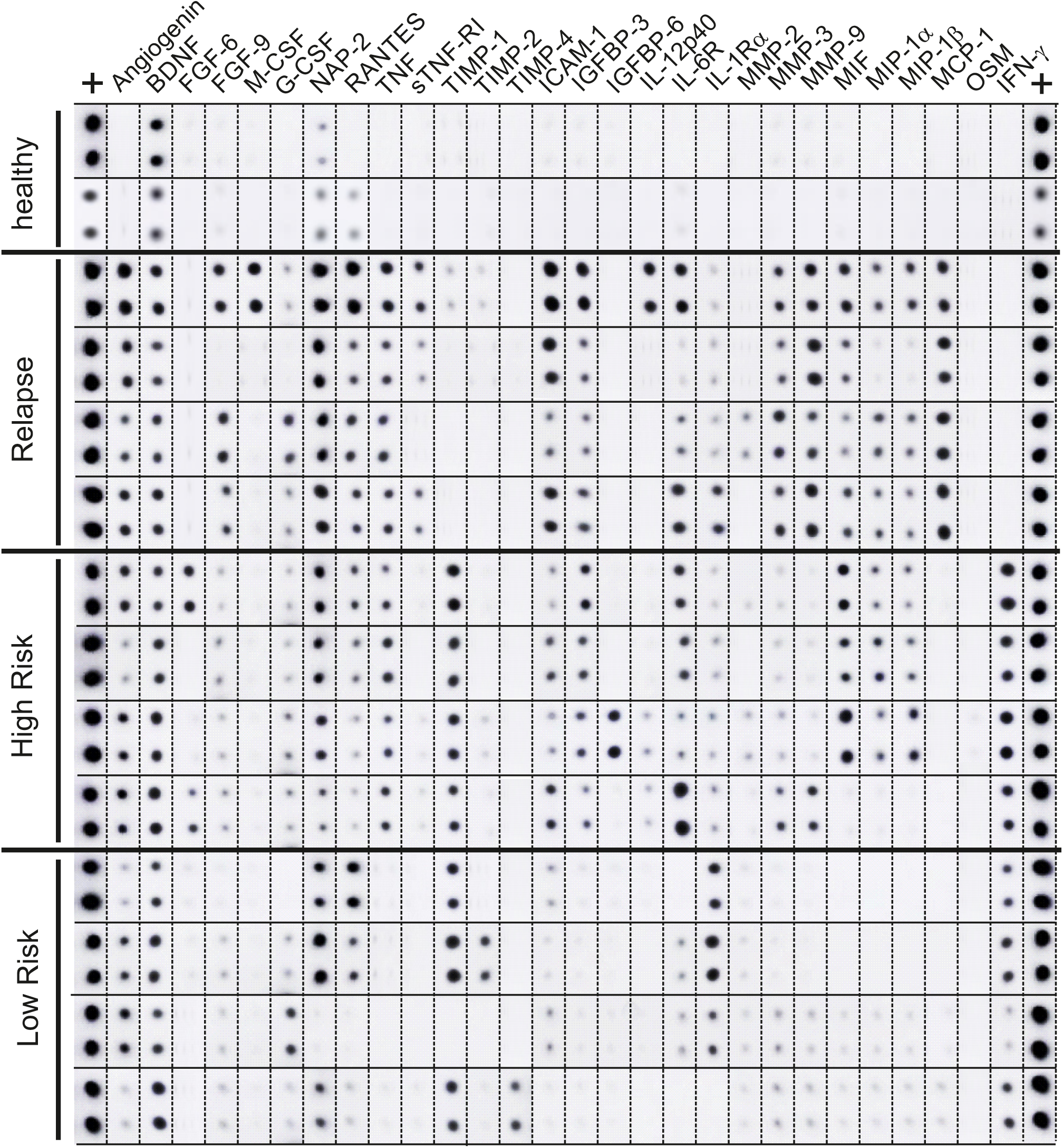
CCF factors detected in pEV from melanoma patients. pEV were purified from 4 ml of plasma by differential centrifugation. The total yield from 1 ml was analyzed for the presence of CCF by a commercial array (Ray Biotech) for 12 patients in three different clinical stages as explained in the text and two healthy controls. Shown are the original signals which were rearranged for better overview and comparison. The positive control (first and last row) is an internal control of the array confirming and measuring the reactivity of the secondary antibody, used to detect the specific signals. This internal control was used to calculate the relative pixel density of the signal (see also **Fig 5A** and **B**).

**Table S1:** Assignment between categories and their reference categories for the overrepresentation analysis. Each patient- and culture category was separated into 3 reference groups: patient-derived pEV, in vitro monocultures and in vitro co-cultures (upper part of Table). In each of these groups independently, miRNAs were analyzed for overrepresentation by checking if the normalized expression value (see details in material and methods) exceeded the highest value from the reference categories (lower part of Table) by at least two-fold. MiRNAs for which this test turned out positive were deemed to be associated with the corresponding category. The overlap of associated miRNAs between selected categories was then inspected using Venn diagrams (**Figure 2B**), and imbalances in the distribution were checked on overlap-derived contingency tables with the chi-squared test (**Figure 2C**).

| Reference group | Sample sources |
| --- | --- |
| Patient-derived blood plasma | healthy, relapse, high-risk, low-risk |
| *In vitro* monocultures | Huh7, Lx2, SkHep1, Tumor, PBMC |
| *In vitro* co-cultures | Huh7-Tumor, Lx2-Tumor, SkHep1-Tumor, PBCM-Tumor |

**Table S2:**
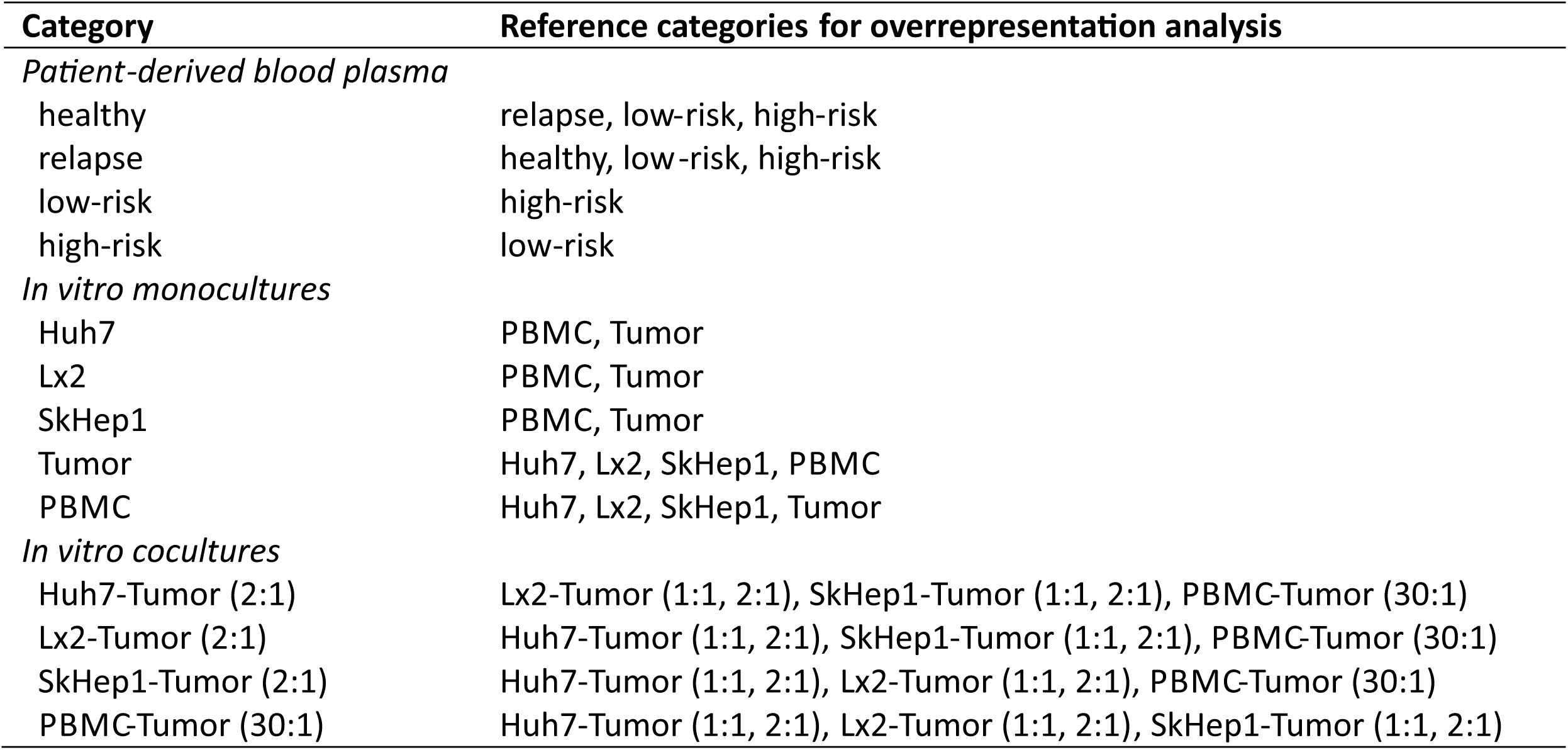
List of miRNAs found in patients pEV and liver/PBMC-melanoma co-cultures. The miRNA lists correspond to the numbers depicted in the Venn diagrams in **Fig 2B** and represent those found in pEV of the respective patient group (relapse, high risk, low risk) and in co-cultures of either PBMC or liver cells with 3 melanoma cells (overlap). Only miRNAs are listed that were at least two-fold higher than in reference groups as defined in **Table S1**.

**Table S3:** Clinical details of patients (cohort 2) from whom plasma samples were analyzed. List of patients (cohort 2) and their clinical parameters who’s plasma samples were used/analyzed in the experiments described in **Fig 2** and **5**. m=male; fe=female.

## Table S02

Representative pEV/EV miRNAs in patient groups (plasma), cell mono-cultures and co-cultures. Only miRNAs are listed that were at least two-fold higher than in reference groups as defined in Table S1 and displayed in the Venn Diagram in **Fig 2B**.

### 1) patients pEV and PBMC

#### Re-patients

miR-382-5p miR-369-3p miR-378i miR-378f miR-1197 miR-197-5p miR-378g miR-1295a miR-487a-3p miR-154-5p miR-3185 miR-346 miR-1306-3p miR-548ad-3p

#### HR-patients

miR-1972 miR-575 miR-140-3p miR-149-5p miR-3180 miR-651-5p miR-6724-5p miR-1236-3p miR-561-5 miR-193a-3p miR-3182 miR-92b-3p miR-155-5p miR-1193 miR-217 miR-1304-5p miR-654-5p miR-592 miR-889-3p miR-1253 miR-548al miR-1246 miR-450b-3p miR-1827 miR-603 miR-381-3p miR-4536-5p miR-2682-5p miR-1273c miR-3180-3p miR-376a-2-5p miR-194-5p miR-422a miR-320e miR-130b-3p miR-4516 miR-363-3p miR-3065-5p miR-494-3p

#### LR-patients

miR-503-5p miR-30c-5p miR-29a-3p miR-340-5p miR-424-5p miR-324-5p miR-518b miR-363-5p miR-151a-3p miR-374b-5p miR-98-5p miR-24-3p miR-19b-3p miR-29b-3p miR-20a-5p+hsa-miR-20b-5p miR-221-3p miR-376a-3plet-7d-5p miR-1183 miR-29c-3p miR-32-5p miR-425-5p miR-431-5p miR-151a-5p miR-23b-3p miR-30e-3p miR-922 miR-18a-5 pmiR-598-3 pmiR-301a-3 pmiR-4286 miR-421 miR-551b-3p miR-10a-5p miR-301a-5p miR-186-5p miR-1255a miR-30e-5p miR-335-5p miR-1537-3p miR-664a-3p miR-337-3p miR-199a-5p miR-454-3p miR-27b-3p miR-625-5p miR-136-5p miR-222-3p miR-323a-3p miR-146a-5p miR-376c-3p miR-22-3p miR-106a-5p+hsa-miR-17-5p miR-148b-3p miR-219b-3p miR-495-3p miR-33a-5p miR-661 miR-486-3p let-7f-5p miR-361-5p miR-30d-5p miR-99b-5p miR-1286

#### PBMC

miR-320a miR-3934-5p miR-1293 miR-606 miR-342-3p miR-132-3p miR-3605-5p miR-650 miR-1973 miR-328-5p miR-210-5p miR-498 miR-1224-3p miR-3615 miR-223-3p miR-520b

#### PBMC-melanoma cell co-culture

miR-1257 miR-579-5p miR-212-3p miR-302b-3p miR-2053 miR-620 miR-644a miR-339-5p miR-142-3p miR-944 miR-548a-5p miR-522-3p miR-1305 miR-548y miR-888-5p miR-627-3p miR-146b-5p miR-506-5p miR-502-3p miR-605-5p miR-3151-5p

#### HR patients & PBMC

miR-181b-2-3p

#### Re patients & LR patients

miR-499a-5p miR-543 miR-769-5p miR-377-3p

#### LR patients & PBMC-melanoma co-culture

miR-1290 miR-107

#### HR patients & PBMC-melanoma cell co-culture

miR-656-3p miR-96-5p miR-520d-3p

#### PBMC-melanoma cell co-culture & PBMC

miR-150-5p

#### LR patients & PBMC

miR-26b-5p miR-361-3p miR-652-3p

### 2) patients pEV and liver cells

#### Re-Patients

miR-382-5p miR-369-3p miR-378imiR-3185

#### HR Patients

miR-575 miR-181b-2-3p miR-140-3p miR-149-5p miR-651-5p miR-6724-5p miR-193a-3p miR-3182 miR-92b-3p miR-155-5p miR-1193 miR-217 miR-1304-5p miR-520d-3p miR-654-5p miR-592 miR-889-3p miR-548al miR-450b-3p miR-1827 miR-603 miR-381-3p miR-4536-5p miR-1273c miR-3180-3p miR-376a-2-5p miR-320e miR-130b-3p miR-4516 miR-363-3p

#### LR Patients

miR-30c-5p miR-29a-3p miR-518b miR-363-5p miR-151a-3p miR-98-5p miR-24-3p miR-19b-3p miR-29b-3p miR-20a-5p+hsa-miR-20b-5plet-7d-5p miR-29c-3p miR-32-5p miR-425-5p miR-431-5p miR-151a-5p miR-23b-3p miR-30e-3p miR-18a-5p miR-301a-3p miR-26b-5p miR-421 miR-107 miR-30e-5p miR-335-5p miR-1537-3p miR-664a-3p miR-454-3p miR-323a-3p miR-146a-5p miR-376c-3p miR-361-3p miR-106a-5p+hsa-miR-17-5p miR-148b-3p miR-652-3p miR-495-3p miR-33a-5p miR-486-3p miR-1286

#### liver cells (huh7, Lx-2, SkHep)

miR-574-5p miR-188-5p miR-302b-3p miR-608 miR-2053 miR-302d-3p miR-215-5p miR-379-5p miR-423-3p miR-518f-3p miR-329-5p miR-548ah-5p miR-325 miR-3144-3p miR-595 miR-339-5p miR-449c-5p miR-483-3p miR-891b miR-1287-5p miR-548ar-5p miR-640 miR-423-5p miR-499b-3p miR-509-3-5p miR-4787-5p miR-216a-5p miR-520f-3p miR-3136-5p miR-323b-3p miR-1285-3p miR-526a+hsa-miR-518c-5p+hsamiR-518d-5p miR-1299 miR-491-5p miR-219a-5p miR-601 miR-612 miR-548a-5p miR-4284 miR-455-5p miR-200a-3p miR-522-3p miR-548e-5p miR-1285-5p miR-183-5p miR-208b-3p miR-299-5p miR-449b-5p miR-4707-5p miR-548y miR-199b-5p miR-627-5p miR-939-5p miR-301b-3p miR-1268b miR-548ar-3p miR-301b-5p miR-548m miR-548v miR-3613-3p miR-3168 miR-885-5p miR-18b-5p miR-582-3p miR-4792 miR-106b-5p miR-4425 miR-197-3p miR-1270 miR-514b-3p miR-205-5p miR-520h miR-506-5p miR-502-3p miR-1178-3p miR-573 miR-663a miR-641 miR-92a-1-5 pmiR-1269b miR-1257 miR-4755-5p miR-1269a miR-138-5p miR-212-3p miR-584-5p miR-192-5p miR-5196-5p miR-101-3p miR-4443 miR-885-3p miR-1268a miR-6721-5p miR-4435 miR-551a miR-128-1-5p miR-145-5p miR-34c-5p miR-654-3p miR-518d-3p miR-626 miR-5001-5p miR-30a-3p miR-584-3p miR-3147 miR-4461 miR-615-3p miR-370-3p miR-181a-3p miR-134-3p miR-450a-2-3p miR-1272 miR-210-3p miR-10b-5p miR-542-3p miR-887-5p miR-597-5p miR-495-5p miR-137 miR-330-3p miR-1297 miR-3131 miR-605-5p miR-1185-1-3p miR-566 miR-129-2-3p miR-620 miR-433-5p miR-616-3p miR-190a-5p miR-1233-3p miR-28-3p miR-769-3p miR-324-3p miR-25-5p miR-95-3p miR-891a-5p miR-133a-3p miR-378d miR-1245a miR-556-5p miR-410-3p miR-1245b-5p miR-224-5p miR-298 miR-936 miR-4707-3p miR-765 miR-433-3p miR-4455 miR-629-5p

#### Liver cells - melanoma cell co-culture

miR-4451 miR-122-5p miR-579-3p miR-143-3p miR-451a miR-381-5p miR-580-3p miR-563 miR-660-3p miR-5010-3p miR-378e miR-513a-3p miR-631 miR-100-5p let-7i-5p let-7b-5p miR-30b-5p miR-190a-3p miR-21-5p miR-190b miR-218-5p miR-92a-3p miR-596 miR-193a-5p+hsa-miR-193b-5 pmiR-3195 miR-132-3plet-7g-5p miR-767-5p miR-296-5p miR-125a-5p miR-9-5p miR-337-5p miR-411-5p miR-1304-3p

#### Re patients & Liver cells

miR-378f miR-1197 miR-1295a miR-346 miR-1306-3p miR-548ad-3p miR-197-5p miR-154-5p

#### HR patients & liver cell-melanoma cell co-culture

miR-1253 miR-1246 miR-2682-5p miR-194-5p miR-422a miR-3065-5p miR-494-3p

#### LR patients & Re patients

miR-499a-5p miR-377-3p

#### LR patients & liver cells

miR-374b-5p miR-376a-3p miR-1183 miR-922 miR-551b-3p miR-1290 miR-301a-5p miR-186-5p miR-1255a miR-337-3p miR-199a-5p miR-625-5p miR-136-5p miR-219b-3p miR-661 miR-99b-5p

#### HR patients & liver cells

miR-656-3p miR-1972 miR-96-5p miR-3180 miR-1236-3p miR-561-5p

#### liver cells & liver cell-melanoma cell co-culture

miR-1296-3p miR-642a-3p miR-564 miR-450a-5p miR-185-5p miR-30a-5p miR-515-5p miR-181c-5p

#### LR patients & liver ceII-melanoma cell co-culture

miR-340-5p miR-424-5p miR-324-5p miR-221-3p miR-598-3p miR-4286 miR-27b-3p miR-222-3p miR-22-3p let-7f-5p miR-361-5p miR-30d-5p

#### Re patients & liver cell-melanoma cell co-culture

miR-487a-3p

#### Re-patients & liver cells & liver cell-melanoma cell co-culture

miR-378g

#### LR patients & Re patients & liver cells

miR-543

#### LR patients & liver cells & liver cell-melanoma cell co-culture

miR-503-5p miR-10a-5p

#### LR patients & Re patients & liver cells & liver cell-melanoma cell co-culture

miR-769-5p

